# Hierarchical neural integration of musical structure during expert performance

**DOI:** 10.64898/2026.07.20.738980

**Authors:** Jamal A. Williams, Coraline Rinn Iordan, Riesa Y. Cassano-Coleman, Elise A. Piazza

## Abstract

Music, which is organized hierarchically (notes, phrases, sections), provides an ideal model for studying fine-grained motor sequence production. However, it is unknown how musicians’ brains integrate tonal structure over multiple timescales during real-life performance. Here, we scrambled an unfamiliar Tchaikovsky piano suite at four timescales—every 1/2/8 measures, or fully intact—and asked expert pianists to play (sightread) all four versions in the fMRI scanner. Responses in the motor network, default mode network, and hippocampus were strongly impacted by scrambling, indicating that they integrate tonal structure over relatively long timescales. Additionally, the emergence of functionally connected sub-networks between auditory, visual, motor, and default mode network regions across scramble levels supported this hierarchical integration process. Our results cannot be explained by lower-level cues (tempo, timbre, dynamics; local pitch height or rhythmic density) and instead reflect processing of high-level tonal structure. Our study highlights novel mechanisms of complex auditory-motor action planning during live music performance.

## Introduction

To complete everyday actions like making a sandwich, crossing the street, and hitting a tennis ball, humans must produce complex motor sequences. Among all possible domains of motor production, musical performance requires processing and planning information that is uniquely hierarchical: music is (precisely and often recursively) organized into nested sections, phrases, and sub-phrases, and listeners develop a sense of musical syntax (e.g., tonal rules about how harmonic sequences should logically progress) both through formal training and implicit exposure (Krumhansl, 2001; Cassano-Coleman, Izen, & Piazza, 2026). Much research has investigated the neural underpinnings of this process for music perception (e.g., Koelsch et al., 2013). However, very little is known about how the brain represents this hierarchical tonal structure during music production, primarily due to the challenge of asking performers to play complex chordal music in the context of a neuroimaging experiment. Here, we addressed this question by systematically scrambling (temporally shuffling) naturalistic piano music at several timescales (measure, sub-phrase, phrase) and asking expert pianists to play (sightread) these different versions on a realistic, three-octave keyboard in the fMRI scanner. This manipulation allowed us to investigate the hierarchical tuning of a broad network of auditory, motor, and higher-order (default mode network, or DMN) regions to these different timescales of musical structure, as well as how these structural shifts impact functional connectivity between these regions. Importantly, our design specifically manipulated tonal structure–a cognitive schema based on higher-order relationships between notes and harmonies within a key context (Krumhansl, 2001)–to reveal a novel neural timescale hierarchy for processing high-level musical information.

In terms of music perception, many studies have investigated the effects of manipulating different aspects of musical structure on the brain. One line of research scrambled recordings of music at a single, short timescale (∼300 ms; Levitin & Menon, 2003; Abrams et al., 2011; 2013) to test which brain regions represent the acoustic structure of natural music (vs. spectrally and temporally matched controls). Relatedly, Fedorenko et al. (2012) probed neural processing of musical pitch vs. rhythm by randomly jittering individual notes on each of those dimensions. A few other studies have focused on hierarchical processing during perception by comparing multiple timescales at once. For example, Koelsch et al. (2013) found that both musicians’ and non-musicians’ EEG responses reflected sensitivity to disruption of long-distance syntactic dependencies in Bach chorales, even when local syntactic structure remained intact. Furthermore, a recent study found that certain regions of the DMN (e.g., medial prefrontal cortex) represent longer-timescale musical events than other DMN regions (precuneus, angular gyrus) and auditory cortex (Williams et al., 2022). Another study scrambled music at multiple timescales (Farbood et al., 2015), building on parallel work investigating the hierarchical neural processing of speech, at timescales from syllables to paragraphs (Ding et al., 2016; Lerner et al., 2011). However, Farbood et al. (2015) scrambled music at the level of the recording, rather than the composition itself, so neural responses may have reflected lower-level acoustic or performance cues (e.g., timbre, tempo, loudness changes) rather than underlying tonal compositional structure. Here, to eliminate the potential influence of these cues that typically vary in natural performance settings, we scrambled pieces at the compositional level (Figure 1), pianists played them along with a metronome, and live auditory feedback was rendered in MIDI with consistent volume and piano timbre. Further, additional control analyses confirmed that our scrambling manipulation only impacted high-level tonal structure and did not induce local changes in lower-level structural cues (pitch height, rhythmic density).

**Figure 1.**
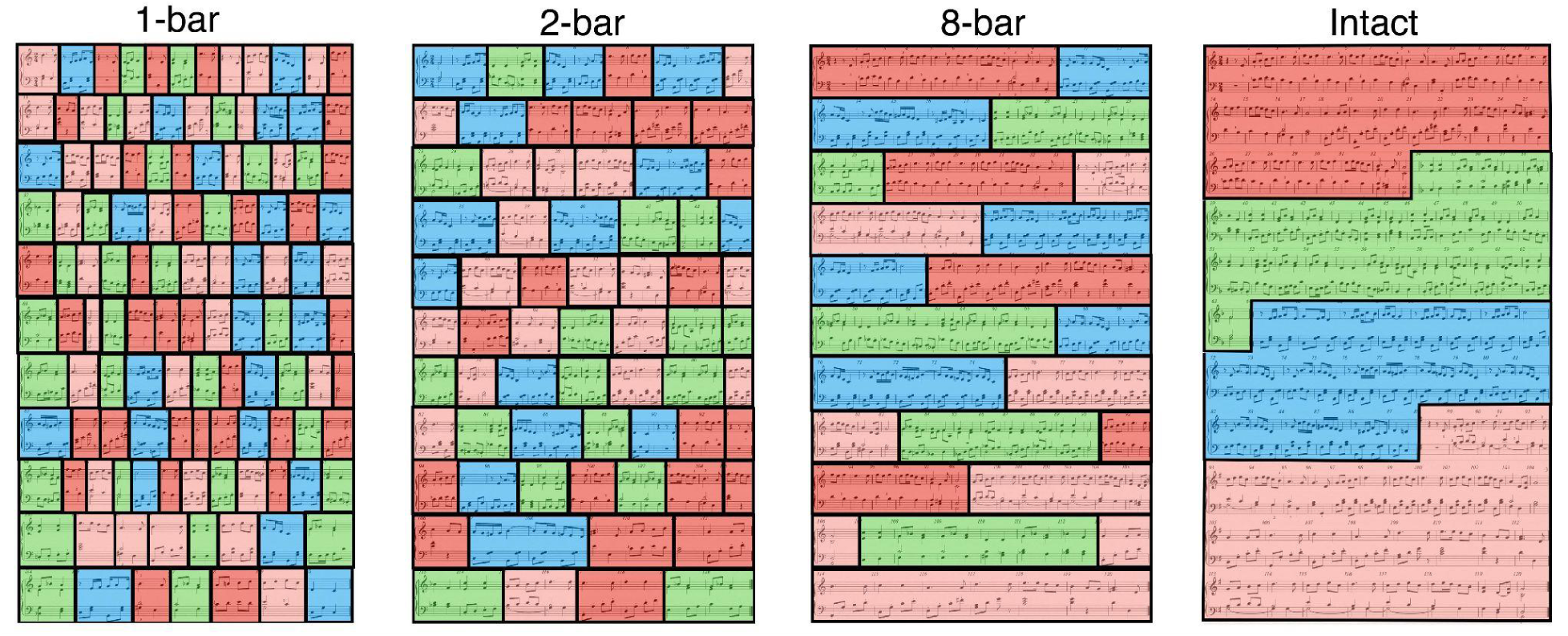
Schematic depiction of musical stimuli in the four scramble conditions. “Intact” shows the full Tchaikovsky medley (adapted from *Album for the Young*), with four sections indicated by color. The other three (scrambled) versions were generated by randomly sampling the Intact version in 1-, 2-, or 8-measure segments. Note that no participants ever saw these colors; they are simply shown here for clarity. In the scanner, during a run of a given condition, participants saw one line (row) of the sheet music at a time, which progressed automatically throughout the run.

Much less is known about hierarchical mechanisms of music production than perception, but one line of fMRI research has focused on rhythmic production, finding that increasing the complexity of rhythmic structure during a simple finger tapping task recruits an extensive network consisting primarily of motor regions (pre-supplementary motor area or pre-SMA, SMA, dorsal premotor cortex, cerebellum, dorsolateral prefrontal cortex), suggesting that this network is involved in hierarchical action planning (Chen et al., 2008a). Chen et al. (2009) propose a theory of auditory-motor interactions whereby sensorimotor transformations occur through relatively indirect connections between superior temporal gyrus and motor cortex via premotor cortex. However, it remains unknown how sensorimotor transformations of pitch-based, tonal structure occur throughout the auditory, motor and higher-order cortices (i.e., DMN regions). We addressed this question by manipulating tonal structure of music via scrambling during naturalistic performance.

A separate line of fMRI research has investigated the production of simple melodic sequences using piano keyboards (or pseudo-instruments) in the scanner, often manipulating access to different modalities of information. For example, Meister et al. (2004) contrasted actual vs. imagined playing of familiar melodies (no auditory feedback in either condition). Other studies have compared passive (motionless) listening with silent playing (no auditory feedback) of either arbitrary notes played on a simple piano keyboard (Bangert et al., 2006), or scales or memorized Mozart pieces performed on a plastic board (Baumann et al., 2007). Pfordresher et al. (2014) compared different kinds of auditory feedback (i.e., timing delays vs. altered pitches) during production of simple melodies, finding dissociable neural networks supporting the integration of perception and action at the timescales of note onset synchrony versus sequential pitch processing. However, no studies have systematically manipulated the tonal syntax of naturalistic music during expert performance on a realistic instrument.

Here, we investigated expert performance on a real, 3-octave keyboard (which enabled the manipulation of complex, hierarchical harmonic structure). We were also specifically interested in the influence of auditory feedback on widespread neural processing of this structure, so we contrasted two different performance conditions: hearing oneself play vs. mute playing (both conditions with sheet music and an audible metronome). We predicted that auditory feedback (which provides significantly more information about nested tonal structure than the sheet music alone) would strengthen the effects of scrambling in auditory regions, but we were also curious about how it might impact motor and higher-order DMN regions, as well as connectivity between these regions.

To summarize, despite extensive previous work on hierarchical mechanisms of music perception, very little is known about such mechanisms during music production, which offers a rich, real-life opportunity to understand complex motor sequence production more broadly. Although a few studies have manipulated rhythmic complexity during production, none have tested how complex tonal structure (reflecting the schematic build-up of structural relationships between pitches and chords) becomes hierarchically integrated across sensory, motor, and default mode network regions during live performance. Here, we asked expert pianists to play naturalistic chordal music–which had been scrambled at multiple timescales (phrases, sub-phrases, single measures with no phrasal structure)--in the fMRI scanner. This scrambled stimulus set was used in a recent behavioral study to test how listeners integrate tonal context across multiple timescales to efficiently predict and remember music (Cassano-Coleman, Izen, & Piazza, 2026).

We focused on two main questions. First, how sensitive are sensory (auditory and visual cortex), motor (motor cortex, cerebellum), and higher-order default mode network regions to different timescales of available tonal information (which our paradigm manipulates via scrambling)? To measure the sensitivity of different regions, we use inter-subject correlation, or ISC (a powerful measure of reliable, stimulus-driven activity that is systematically synchronized across people in a given brain region; Nastase et al., 2019). If we find that a region responds systematically more reliably (higher ISC) for more intact (i.e., less scrambled) music, this suggests that that region integrates tonal structure over increasingly long timescales. Applying this framework, prior work has suggested that the DMN integrates context over relatively long timescales during narrative (Lerner et al., 2011; Baldassano et al., 2017; Yeshurun et al., 2021) and music (Farbood et al., 2015; Williams et al., 2022) perception, whereas primary auditory cortex or A1 (which responds similarly reliably for short-timescale vs. long-timescale scrambling–i.e., word-level vs. paragraph-level; Lerner et al., 2011) represents shorter timescales, but it is unknown how this hierarchical processing extends to production of extended musical sequences. Furthermore, the impact of scrambling tonal structure on motor regions is an even more open-ended question.

Second, how does functional coupling between these regions support this hierarchical integration of tonal context? To test this, we used inter-subject functional correlation (ISFC), a measure of stimulus-driven functional connectivity that depends on intersubject consistency, thus making it less susceptible to intrinsic neural dynamics or non-neuronal artifacts than traditional functional connectivity (Simony et al., 2016; Yeshurun et al., 2021). We predicted that more intact music will strengthen ISFC between auditory and motor regions, as well as between auditory and DMN regions.

Moreover, the potential finding that motor-DMN connectivity may increase with longer-timescale tonal structure would shed new light on how the motor system processes and integrates complex musical content during real-life performance, and provide new mechanisms for the feeling many pianists have of representing and anticipating the music “in their fingers” (Taylor & Witt, 2015; Bernardi et al., 2013). In addition, we exploited the fine-grained naturalistic structure of our stimuli (i.e., sections containing distinct themes) to verify the overall reliability of music-driven neural signals across brain regions. Finally, we conducted several control analyses to verify that the effects of scrambling we report are driven by high-level tonal structure, rather than lower-level musical cues, such as local variation in pitch height or rhythmic density, or variation in performance accuracy.

## Results

### Scrambling reveals hierarchical organization of musical structure in auditory, motor, and higher-order regions during naturalistic music production

To measure the sensitivity of different brain regions to different timescales of tonal structure, we computed intersubject correlation (ISC) in each scramble condition. If a region shows an effect of scramble level (i.e., with systematically higher ISC from the most to least scrambled conditions; Figure 1), that suggests that it integrates information (context) over the increasingly long timescales included in our manipulations (Lerner et al., 2011). If it shows no effect, that suggests it is not sensitive to this additional context. We included a set of regions of interest (ROIs; Figure 2) that we predicted might be particularly involved in tracking temporal tonal structure in the context of music production: primary and secondary auditory cortex (A1, STG), dorsal and ventral premotor cortex, motor cortex, pre-supplementary motor area, supplementary motor area, cerebellum, the default mode network (defined to include angular gyrus, temporal-parietal junction, precuneus, posterior cingulate cortex, and medial prefrontal cortex; Yeshurun et al., 2021), and the hippocampus. We were also interested in testing whether primary visual cortex, potentially involved in representing visual cues from the score that provide information about tonal structure, organized this information hierarchically. We also compared ISC across two conditions: playing with feedback (auditory-motor, or “AM”) and silent playing (“M”).

**Figure 2.**
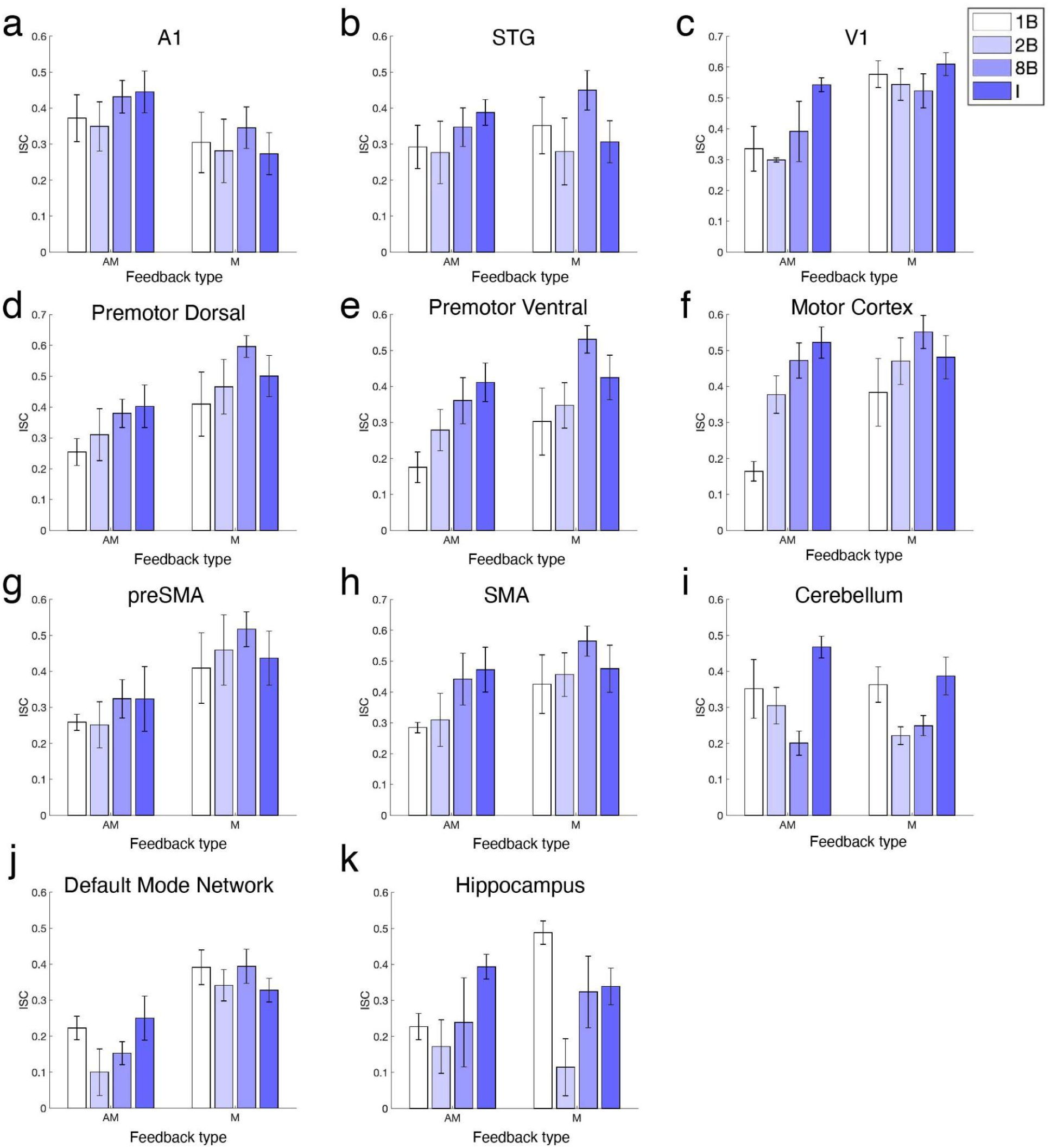
Intersubject correlation (ISC) by region-of-interest and feedback types (“AM” = playing with auditory feedback; “M” = playing without auditory feedback). Error bars are s.e.m.

We conducted a 3-way ANOVA (ROI x scramble condition x feedback condition) and found a significant main effect of scramble condition (*F*(3) = 4.93, *p* < .05), a significant main effect of ROI (*F*(10) = 11.86, *p* < .001), and a significant interaction between ROI and scramble condition (*F*(30) = 5.68, *p* < .001). There was no main effect of feedback (“AM” vs. “M”; *F*(1) = 1.92, *p* = .21), indicating that playing in silence did not impact participants’ overall intersubject reliability in these ROIs as a whole, but a significant interaction between feedback and ROI (*F*(10) = 5.55, *p* < .01) suggested that auditory feedback impacted overall ISC in different ways across regions (e.g., it improved ISC in A1 but dampened it in the DMN). There was no significant interaction between feedback and scramble condition (*F*(3) = 2.04, p = .16), or between feedback, ROI, and scramble condition (*F*(30) = 1.29, *p* = .29), suggesting that in general, auditory feedback did not systematically alter the effect of tonal temporal disruptions on the reliability of these brain regions. All p-values above were adjusted using Greenhouse-Geisser correction.

Because of the significant interaction between ROI and scramble condition, next, we tested more fine-grained effects of scrambling at the individual ROI level. To do so, we ran post-hoc permutation tests comparing the Intact condition with the three other scramble conditions (see Table S1 for full set of contrasts; *p* < .005 after Bonferroni correction). While many ROIs showed an effect of scrambling (reflecting the main effect in the ANOVA above), the precise pattern differed across ROIs, with the premotor and motor cortex showing the most monotonically increasing patterns (Figures 2d-f). In the default mode network and hippocampus (Figures 2j-k), ISC increased systematically between the 2B, 8B, and Intact conditions but was surprisingly high in the most scrambled (1B) condition. We interpret this as an effect of the attentional demands of this extreme level of scrambling, as it is the only condition that fully removes any trace of meaningful phrase structure. This is supported by recent behavioral findings testing listeners’ event perception in this same scrambled stimulus set (Cassano-Coleman, Izen, & Piazza, 2026; Experiment 3): specifically, listeners reliably perceived both 8-bar phrases and 2-bar subphrases, but not 1-bar segments, as “meaningful events”. Thus, in this particularly unstructured, chaotic condition, our participants (expert pianists) may have been highly attentionally engaged in seeking predictable patterns (linking multiple measures) that do not exist, more so than in the other three conditions.

We also conducted a related control analysis to confirm the general reliability of music-related patterns in our neural data, based on correlations between repetitions of the same stimulus (which we call “inter-rep correlation”, or IRC; see Supplementary Information for full details). Specifically, we confirmed that repetitions of the same scramble condition were significantly more correlated than repetitions of different (randomly chosen) scramble conditions (Figure S1 and Table S2). We also computed IRC between repetitions of the Intact condition only, finding that correlations between segments corresponding to the same musical section (colored segments in Figure 1, right; each section contains a cohesive musical theme) were higher than between segments of different (randomly chosen) sections of the piece (Figure S2 and Table S3). Together with our ISC analyses, these results further demonstrate the high reliability of stimulus-driven neural signals generated by this paradigm.

### Networks of auditory, motor, and higher-order brain regions emerge as musicians perform longer-timescale tonal structure

Next, to test how coordination between the regions above may support the hierarchical integration of tonal information, we investigated how scrambling impacts functional connectivity between these regions. Here, we separated the default mode network into individual regions, to visualize how scrambling affected connectivity both within the DMN and between the DMN and other regions. We used intersubject functional correlation (ISFC), a measure of stimulus-dependent inter-regional correlations that are consistent across multiple subjects’ brains and therefore specifically related to stimulus properties rather than to intrinsic neural properties or noise (Simony et al., 2016). Figure 3 shows ISFC matrices (top) and circular connectivity plots (bottom) across the four scramble conditions. Figure S3 depicts significant values after phase-randomization (as in Simony et al., 2016) and correction for multiple comparisons using the Benjamini-Hochberg procedure (q<.05).

**Figure 3.**
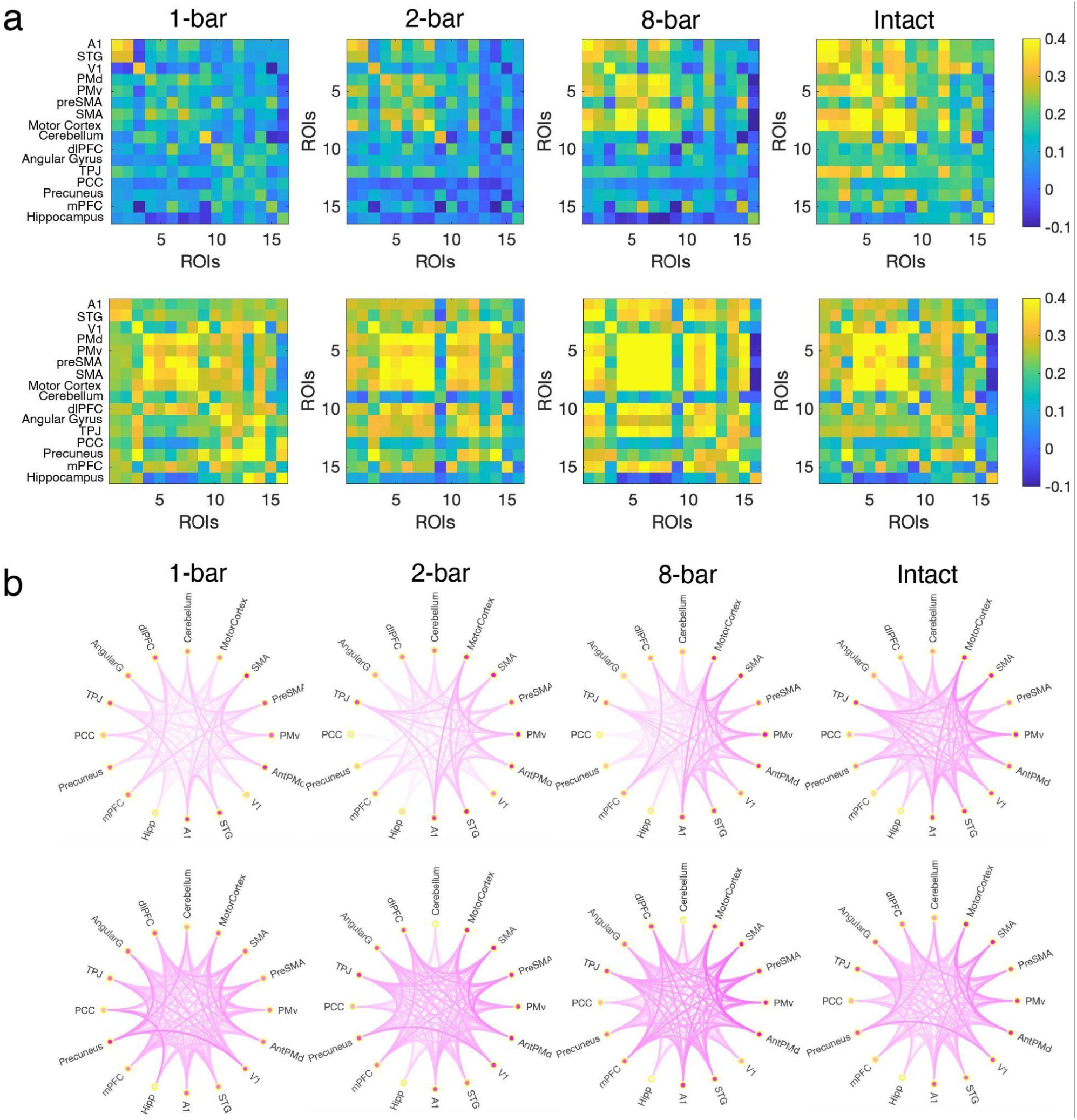
Intersubject functional correlation (ISFC), plotted as (a) correlation matrix and (b) schemaball (circular network graph). Top rows depict “AM” and bottom rows depict “M”. In (b), regions with more saturated (dark orange) nodes have higher average absolute functional correlation with other regions.

In the ISFC maps for “AM” (playing with auditory feedback; Figure 3, top rows), as music becomes increasingly intact (left-to-right), we observed a systematic increase in connectivity between auditory and DMN regions, as well as between these two networks and the motor network and visual cortex. Additionally, we observed increased connectivity within the motor network itself. In “M” (silent playing; Figure 3, bottom rows), ISFC was higher overall, but with less clear systematic effects of scrambling. To assess all of these effects statistically, we first conducted several 2-way ANOVAs (feedback type x scramble condition) for these groupings of ROIs. We found significant main effects of scramble condition on ISFC between A1 and V1 (Figure 4a; *F*(3) = 6.95, *p* < .01), between auditory cortex (A1, STG) and all motor cortical regions (PMd, PMv, preSMA, SMA, motor cortex) (Figure 4b; *F*(3) = 4.71, *p* < .05), within all motor regions (Figure 4d; *F*(3) = 6.85, *p* < .01), between the cerebellum and the default mode network (Figure 4f; *F*(3) = 3.84, *p* < .05; as in ISC analyses, DMN included angular gyrus, temporal-parietal junction, precuneus, posterior cingulate cortex, and medial prefrontal cortex), and between V1 and the DMN (Figure 4g; *F*(3) = 3.90, *p* < .05). We also found interactions between feedback type and scramble level for ISFC between A1 and V1 (Figure 4a; *F*(3) = 8.51, *p* < .01), auditory cortex and the DMN (Figure 4c; *F*(3) = 4.49, *p* < .05), motor cortex and the DMN (Figure 4e; marginally significant: *F*(3) = 3.35, *p* = .08), and V1 and the DMN (Figure 4g; *F*(3) = 4.89, *p* < .01). All p-values above were adjusted using Greenhouse-Geisser correction.

**Figure 4.**
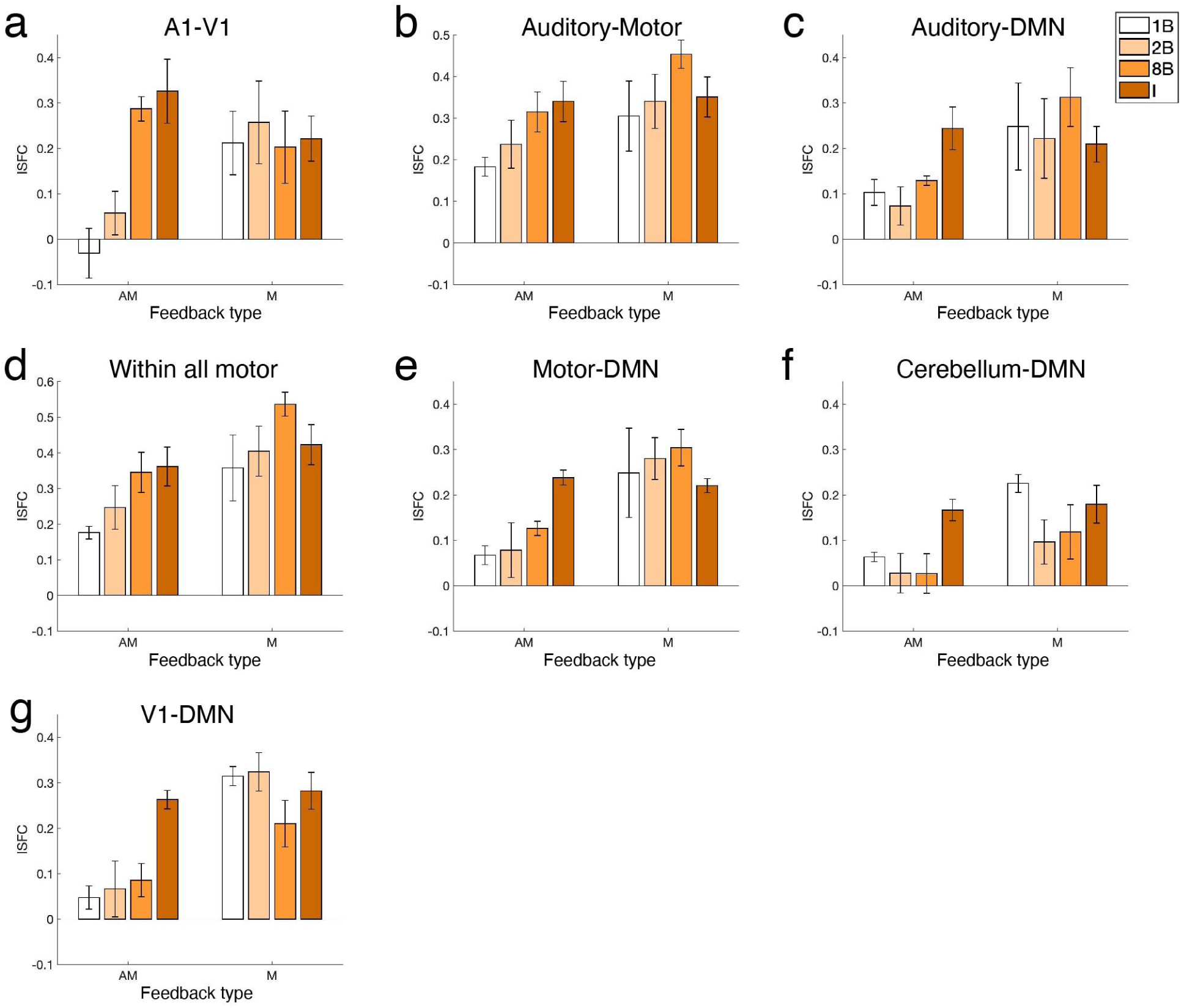
Intersubject functional correlation (ISFC) for different ROI groupings and feedback types (“AM” = playing with auditory feedback; “M” = playing without auditory feedback). Means reflect ISFC (r) values averaged across relevant cells of Figure 3a. Error bars are s.e.m.

Given these interactions, as well as the distinct patterns of scramble effects we observed across ROI groups, we conducted post-hoc permutation tests between the Intact condition and the three scramble levels for “AM” and “M” separately (as we did for ISC; see Table S4 for full set of ISFC contrasts; *p* < .007 after Bonferroni correction).

In general, scrambling tonal information systematically affected connectivity between auditory, visual, motor, and higher-order processing regions, and this effect was generally stronger when players received auditory feedback (“AM”) than when they did not (“M”) (see significant interactions above and individual scramble contrasts in Table S4; no contrasts were significant in the “M” group). In particular, auditory feedback facilitated effects of scrambling on connectivity between A1 and both V1 and the DMN, as well as motor cortex and the DMN, cerebellum and DMN, and V1 and DMN (Figure 4). This strongly suggests that during performance, auditory feedback plays a critical role in bridging hierarchical representations of high-level (tonal) musical structure between regions involved in executing fine motor production and the DMN, which is known to integrate complex structure from naturalistic sounds over long timescales (Yeshurun et al., 2021). In terms of V1-DMN connectivity, the significant feedback type x scramble level interaction was driven by the “Intact” condition, suggesting that auditory feedback selectively boosts the propagation of relatively long-timescale tonal information from visual cortex (e.g., representations of intact phrase structure in the sheet music) to higher-order areas.

### The effects of scrambling on the brain cannot be explained by behavioral (piano-playing) performance or low-level musical cues

Finally, we conducted three additional control analyses to verify that our main results indeed reflected hierarchical processing of tonal structure and could not be explained by potential differences in behavioral performance across conditions or by low-level musical cues.

First, mean performance accuracy did not vary significantly across scramble conditions (Kruskal-Wallis tests, “AM”: *χ*^2^(3) = 1.39, *p* = .71; “M”: *χ*^2^(3) = 2.41, *p* = .49; Figure S4), so it could not explain large scrambling effects (e.g., ISC in motor cortex; Figure 2f).

Second, differing patterns of ISC and ISFC between “AM” and “M” (Figs. 2 & 4) could not be explained by differences in performance accuracy across feedback types, since they did not differ significantly in any scramble condition (Figure S4; see Supplementary Information for additional details).

Third, we sought to rule out the possibility that our scrambling manipulation somehow introduced variability in low-level musical cues at scramble boundaries, which could have theoretically influenced our neural results. Specifically, we measured changes in pitch height (Figure S5a) and rhythmic density (Figure S5b) across scramble boundaries (in the 1B, 2B, and 8B conditions; orange boxes) and found that they were not significantly different than changes in these features across boundaries separating equivalently-sized segments of music in the Intact condition (blue boxes). See Supplementary Information for additional details.

## Discussion

We found novel evidence for hierarchical neural integration of high-level musical syntax during live, complex, expert performance. First, across a broad network of auditory, motor, visual, and higher-order (DMN) brain regions, we report widespread effects of scrambling musical content on the reliability (intersubject correlation) of neural responses, reflecting hierarchical tuning to a range of timescales of naturalistic musical structure (i.e., measure, sub-phrase, phrase). Second, we found that as performers play music containing progressively more intact tonal context, this longer-timescale structure systematically reorganizes functional network structure between auditory, visual, motor, and higher-order regions. This effect is generally stronger when players hear themselves play (“AM” condition) than when they do not (“M” condition), even for connectivity between ROIs outside of auditory cortex (e.g., motor-DMN, cerebellum-DMN, V1-DMN). We also conducted several analyses which verified the overall stimulus-driven reliability of neural responses (i.e., across repetitions of the same musical content). Finally, we confirmed that the effects of scrambling reflect processing of high-level tonal structure: our scrambling manipulation occurred at the compositional level and was rendered in MIDI with no variation in tempo, timbre, or dynamics (loudness), and additional analyses ruled out the influence of lower-level musical cues (i.e., local variation in pitch height or rhythmic density) and variation in performance accuracy. Thus, these neural regions must specifically keep track of high-level tonal information to coordinate the execution of a complex piano piece in expert performers.

These findings contribute significantly to our understanding of everyday motor action planning. While some previous research has focused on manipulating rhythmic complexity during finger-tapping tasks in the scanner, little was known about how the brain integrates hierarchical tonal structure across different timescales during live performance (i.e., which involves integrating contextual information about the linking of several notes into a phrase, two adjacent phrases into a “period”, multiple periods into a section, etc; Knosche et al., 2005; Neuhaus et al., 2006). Some of our results are highly consistent with previous work on finger tapping. For example, as metrical complexity increased, Chen et al., 2008a found increasing recruitment of pre-SMA, SMA, dPMC, and cerebellum; similarly, we found increasingly reliable responses in SMA, dPMC, and motor cortex as performers played increasingly long-timescale tonal structure.

Relatedly, we report an effect of scrambling on auditory-motor connectivity (similar to the effect of metrical saliency reported by Chen et al., 2006), revealing systematic sensorimotor transformation of both rhythmic and harmonic structure. However, Chen et al., 2008b found that as rhythmic complexity increased, activity in dPMC (but not vPMC) increased (in non-musicians doing a finger-tapping task), whereas we found effects of scramble level in both dPMC and vPMC in expert pianists playing complex tonal music. Overall, our findings validate and extend this previously reported motor hierarchy (for simple rhythmic production) to the novel context of production of tonally complex chordal music, and we additionally describe the involvement of auditory, visual, and DMN regions (and connections between them) in this process.

Our study adapted a scrambling paradigm previously used to demonstrate hierarchically organized temporal receptive windows (TRWs; Lerner et al., 2011), analogous to spatial receptive fields but for integrating low-level sensory to higher-order content during naturalistic narrative comprehension. One previous study (Farbood et al., 2015) applied a similar scrambling paradigm to recordings of Brahms and found parallel TRW hierarchies for music vs. speech perception across auditory and DMN regions, but these results could have been influenced by acoustic variability in the stimuli (e.g., large fluctuations in dynamics across the piece). Here, we applied this TRW framework to motor production of complex music scrambled at the level of the composition (rather than the recording) to bring forth a new understanding of hierarchical processing of music. We found systematic effects of scrambling on ISC in regions throughout the motor network (Figure 2), revealing a novel temporal receptive window hierarchy for processing and generating musical syntax during naturalistic performance.

In terms of the patterns of ISC in sensory (A1) and higher-order (DMN) regions, many of our results were highly consistent with prior work on TRWs for listening, but some differed slightly. In previous work (on listening to scrambled speech; Lerner et al., 2011), A1 showed little effect of scrambling, reflecting short-timescale tuning, whereas areas adjacent to A1 along the STG (through pSTS) began to show increasingly longer-timescale tuning. We found a partial effect of scrambling in A1 (specifically, Intact > 2B). It is possible that during active production, feedback connections alter or shift the TRW hierarchy downward; however, in general, the effects of scrambling we report in both A1 and STG are still quite weak overall, consistent with prior work (and, as expected, there is no effect in the “silent motor” condition). In the DMN, we found an expected systematic increase in ISC from “2B” to “Intact”, also consistent with prior TRW work (Lerner et al., 2011; Farbood et al., 2015), but we found surprisingly high ISC in the most finely scrambled (“1B”) condition. This can potentially be explained by the high attentional demands of this condition, resulting in increased engagement which, counterintuitively, increased intersubject alignment (i.e., by generating more highly surprising or salient moments than the other scramble conditions). This explanation fits with an updated definition of the DMN as a dynamic “sense-making” network, supported by extensive literature on narrative processing; Yeshurun et al., 2021). This interpretation is also supported by our recent behavioral finding that the “1B” scramble level is perceived as uniquely unstructured and random (i.e., listeners fail to reliably segment this condition into coherent events, unlike 2B and 8B; Cassano-Coleman et al., 2026). Another interesting finding is that the DMN’s connectivity (with auditory, visual, and motor regions) dramatically decreases with increased scrambling (Figure 4c, 4e-g). This may signify a distinction between a more connected network structure during “typical task mode” (i.e., the “Intact” condition with auditory feedback, in which musical structure contains long-timescale structure consistent with the DMN’s well-learned schemas), and increasingly scrambled conditions, in which the DMN begins to decouple from these other systems to rely on a more internalized pattern-seeking or “sense-making” mode (Yeshurun et al., 2021). Overall, these findings invite further investigations of the DMN’s role in dynamic processing, integration, and memory for music, especially given the distinct structure of naturalistic music vs. narratives (Besson & Schon, 2001; Temperley, 2022).

Our design contrasted music performance with and without live auditory feedback (“AM” vs. “M”). Interestingly, we found no main effect of feedback type on ISC, indicating it did not impact the overall reliability of neural responses across ROIs, and no interaction between feedback type and scramble condition, or feedback type, ROI, and scramble condition, indicating that effects of scrambling were not reliably differentiable between these two conditions. However, there were differential effects of feedback on overall ISC within specific ROIs (indicated by a significant interaction between feedback type and ROI). In A1, ISC was stronger overall in “AM” than “M”, while in the DMN and V1, it was the reverse. This is consistent with the interpretation that auditory information should increase the reliability of auditory responses, whereas in silence, with fewer cues to musical structure available, participants may be more engaged in internally guided thinking, or more focused on salient visual markers in the sheet music. Motor regions showed less of an impact of auditory feedback than the other ROIs, as expected, since motor output was the same regardless of feedback. However, auditory feedback did strengthen the effects of scrambling on connectivity between the DMN and auditory, visual, and motor regions (Figure 4), suggesting that during performance, auditory feedback connects hierarchical representations of musical syntax across sensory, motor, and higher-order regions.

Our results also complement related fMRI work on musical imagery; for example, Meister et al. (2004) compared pianists’ actual vs. imagined performance of familiar (recently practiced) right-hand melodies (no auditory feedback in either condition). They found some shared activation of the precuneus, intraparietal sulci, and visual cortex in both conditions, whereas the left primary sensorimotor area and the cerebellum were selectively active during silent playing (not imagery). This dissociation provides some evidence that auditory imagery is unlikely to explain our robust scrambling effects in the “silent playing” group. Furthermore, we verified that our stimuli were unfamiliar to all participants, and our “M” group never played the stimuli with auditory feedback at any point, making it unlikely that imagery played a strong role. Our participants also had surprisingly weak explicit representations of the high-level structure of the stimuli: when asked after the session how they thought the pieces were constructed, they were all unaware of the manipulation (i.e., none guessed that they were playing a real piece that had been scrambled; see Methods). Still, future work directly comparing a more comprehensive set of tasks could further disentangle the mechanisms of imagined, silent, and naturalistic playing.

By combining the manipulation of fine-grained structure of music during live, naturalistic performance on a realistic piano keyboard in the scanner, our paradigm opens new possibilities for future investigations of real-life music production. For example, future studies could disentangle hierarchical mechanisms of pitch vs. rhythmic structure in live performance; some prior work has independently manipulated these dimensions in the context of simple melodic production (Bengtsson & Ullen, 2006), but it is less known how multiple hierarchical levels of pitch and rhythmic complexity (e.g., hypermeter, polymeter and other complex forms) become integrated during performance of polyphonic music, and coordinated across musicians in ensembles.

Further, building on prior neuroimaging work comparing high-level structural vs. low-level motor planning (i.e., specific finger movements; Bianco et al., 2022), it will be important to investigate how the brain integrates hierarchical tonal structure in a generalizable way across different low-level instantiations (fingerings, hand configurations that vary across instruments).

The influence of musical learning and expertise on mechanisms of motor production is also a relatively open area, and certainly challenging to test causally; one longitudinal study tested non-musicians’ learning of simple sequences on the cello (Wollman et al., 2018), finding increased activation in the dorsal auditory network and increased functional connectivity between the superior parietal lobule, supplementary motor area, and auditory cortex. Olszewska et al. (2024) found that over the course of piano training, there was less involvement of higher-order cognitive control and integrative regions, as well as of the basal ganglia, but it was difficult to disentangle effects of training and piece difficulty. Given that our participants were highly trained (mostly professional; a few semi-professional) pianists, future work incorporating manipulations of musical structure into longitudinal musical training paradigms–including with young children–would shed light on how the hierarchical mechanisms we report develop over the course of learning a specific instrument, and how they relate to explicit knowledge of Western tonal theory. There has been some work exploring mechanisms of music improvisation, including in interacting pairs (Donnay et al., 2014); such studies have mainly focused on mechanisms relating to reward, creativity and self-monitoring (Limb & Braun, 2008; McPherson et al., 2016; Barrett et al., 2025), as well as interbrain synchronization (Mueller et al., 2013), but further research is needed to determine how these processes relate to fine-grained representations of sequential action patterns generated by oneself vs. others (e.g., Kohler et al., 2023) during complex musical interactions.

Our study reveals new mechanisms of hierarchical musical structure integration during naturalistic performance. As expert performers played music that contained progressively longer-timescale intact tonal structure in the fMRI scanner, we found systematic increases in the stimulus-driven reliability (measured with intersubject correlation) of auditory, motor, and higher-order processing regions, indicating that these regions integrate relatively long-timescale tonal information. We also found systematic increases in coupling between auditory, visual, motor, and default mode network regions (measured with intersubject functional correlation), indicating the emergence of functionally connected networks to support hierarchical, action-based musical phrase processing and clarifying the role of the DMN in auditory-motor sequencing. Our design controlled for low-level acoustic cues (dynamics, timbre, tempo), and several additional analyses ruled out the influence of local musical cues (pitch height, rhythmic density) or variation in performance accuracy, ensuring that our effects are due to high-level tonal structure. Future research could investigate how this hierarchical neural integration changes across the lifespan, including in disorders like Parkinson’s and Alzheimer’s disease, potentially leading to more effective and tailored music-based therapeutic approaches.

## Methods

### Participants

Nine right-handed, healthy participants (3 females, ages 19-38; mean = 26.1 yrs, SD = 6.0 yrs) completed the study. All were highly proficient pianists recruited from a network of professional piano teachers and performers in the Princeton, NJ area, as well as the Princeton Pianists’ Ensemble. None of the subjects had any history of neurologic, auditory, or psychiatric disorders. Informed consent was obtained in writing for all subjects, and the research protocol was approved by the Institutional Review Board at Princeton University. Participants were paid $20 per hour. Six additional participants were excluded because they did not achieve sufficient performance accuracy levels in either the mock scanner pre-test (N=4) or the real fMRI scan (N=2). Four participants completed the “AM” (audio-motor) condition and five participants completed the “M” (silent motor) condition (random assignment of groups). We chose this between-subjects design to eliminate contamination between feedback conditions.

Recruiting eligible participants was particularly difficult, given that our task (sight-reading complex, unfamiliar, scrambled chordal music inside the scanner, with a high degree of accuracy) was extremely challenging compared to related previous studies that primarily used simple melodic sequences. The only participants who met our performance accuracy standards for this task were professional or semi-professional pianists (performers and/or teachers), with extensive experience as accompanists for musicals, choirs, etc. Despite these challenges, our sample size was nevertheless highly comparable to other studies that required people to play musical instruments in the fMRI scanner (Wollman et al., 2018; Donnay et al., 2014; Limb & Braun, 2008; Bangert et al., 2006; Bengtsson & Ullen, 2006; Meister et al., 2004). Because of these challenges related to sample size, we took additional measures to rigorously test the reliability of our data, including within-subject control analyses that tested our ability to measure reliable neural representations of fine-grained musical content (i.e., comparing runs of the same scramble condition vs. random conditions (Figure S1) and comparing repetitions of the same section of the Intact condition vs. random sections (Figure S2); see Supplementary Information for details).

### Musical stimuli

To generate the “Intact” musical piece subjects performed in the scanner (120 measures in 2/4 time, 60 bpm, 4 min total; Figure 1, panel 4), we sampled several sections from Tchaikovsky’s piano suite *Album for the Young*, which was chosen because it contained complex harmonic and melodic structure but would still be relatively easy for experts to sight-read under the challenging constraints of the fMRI scanner environment. Every measure was 2-s long (1 beat = 1 s). To create the scrambled versions of this piece (Figure 1, panels 1-3), we temporally shuffled it at several timescales: 8-bar (i.e., 8-measure segments of intact context), 2-bar, and 1-bar. No segments were contiguous from the original music. Unlike previous studies testing the influence of dynamic musical structure on the brain (Williams et al., 2022; Burunat et al., 2024), including those using scrambled music (Levitin & Menon, 2003; Abrams et al., 2011; Farbood et al., 2015), here, we scrambled the composition itself (rather than the recording), pianists played along with a metronome, and live auditory feedback was rendered with uniform timbre and volume (see Procedure below), thus fully eliminating the influence of these cues and ensuring that we were specifically disrupting tonal structure.

### Mock scanner session

Before the MRI scan session (on a separate day), all participants completed a behavioral pre-test in the MRI simulator, or “mock scanner” (an environment designed to simulate the feel and sounds of being in an actual scanner), using the same keyboard as in the real scan. The goal of this was to familiarize participants with the piano performance task within the constraints of the scanner environment and to ensure that they could perform sufficiently accurately (∼75% accurate notes). The tasks were identical to those of the main scan session. However, stimuli performed in the mock scanner pre-test were sampled from *different* sections of Album for the Young (to give participants practice playing the same general style of music without pre-exposing them to the stimuli used in the actual scan). We used the same procedure as above to generate new pieces with four levels of scrambling (Intact, 8B, 2B, 1B).

### Procedure and piano keyboard setup

In the main fMRI scan session, all participants completed twelve, four-minute functional runs of piano playing (three identical repetitions of each of the four scramble conditions above). Conditions were pseudo-randomized, such that all subjects first played all four scramble conditions within a first “block”, then played them again in a second “block”, then a third “block”, with the order of conditions within a block counterbalanced across blocks and across participants, with no contiguous repetitions of any conditions. In each run, sheet music for that condition was presented via back projection on a screen placed at the end of the MRI bore, one line of music at a time (to ensure alignment of visual stimuli across subjects). To additionally ensure that participants’ playing would be aligned across runs, a metronome (sine-wave beep at 60 bpm) played during every run, including a 2-measure count-off (which subjects practiced beforehand), and this sine-wave was time-locked with the scanner trigger via MATLAB. The (3-octave) keyboard was custom-built by a Masters candidate in Medical Biophysics at National Taiwan University to be MRI-compatible (with plastic keys and casing), and its audio output was transmitted to a MacBook Air laptop in the control room and recorded with GarageBand (Apple Inc., Cupertino, CA). For subjects in the “AM” group, live (imperceptibly low-latency) auditory feedback was played back to them diotically over MRI-compatible earbuds (Sensimetrics corp.) at a comfortable listening level that could be easily heard over the background scanner noise while they played each piece (i.e., live MIDI feedback was rendered in GarageBand using a natural grand piano timbre). Subjects in the “M” group played silently with no auditory feedback (except for the metronome). MRI-compatible earplug putty (Mack’s corp) was molded around the Sensimetric earbuds to hold them in place and offer additional noise protection. To minimize motion during this unique task, we 3D-printed foam head cases (produced by Caseforge; https://caseforge.co) based on models of each participant’s head built during their initial (mock scanner) visit. These head cases were uncomfortable for some participants, so as an alternative we used a combination of medical tape across the forehead and MRI-compatible cushions placed between the participant’s ears and the edges of the head coil, which has been shown to be highly effective in naturalistic tasks (Jolly, Sadhukha, & Chang, 2020). Overall, these methods were very successful in minimizing head motion (Figure S6). The piano keyboard rested on each subject’s lap in supine position, with their knees elevated by a large bolster cushion. Between runs, subjects were allowed to take short breaks inside the scanner (i.e., closing their eyes and resting their hands). At the start of the session, participants were given 5 minutes to improvise freely on the keyboard (no sheet music) with the scanner off, to test the auditory feedback and behavioral recording and make any necessary adjustments to their body position and cushions to optimize their comfort. After completing all playing runs and exiting the scanner, we asked all participants to guess how the musical stimuli were constructed, and none guessed correctly (i.e., none mentioned scrambling, shuffling, or reordering, but 1-2 thought that they were designed to “sound weird” or were simply “poor quality compositions written by scientists”).

### Data acquisition and preprocessing parameters

Scanning sessions took place at the Scully Brain Imaging Center at the Princeton Neuroscience Institute on a whole-body 3T MAGNETOM Prisma MR scanner with a 64-channel head coil. Each scan session included one sagittal T_1_-weighted image (MPRAGE, voxel size 1 mm^3^) for anatomical reference. We recorded echo-planar imaging (EPI) images covering the whole head (voxel size 3 mm^3^, 50 interleaved slices, echo time (TE) 30 ms, TR 1700 ms, flip angle = 70°).

Results included in this manuscript come from preprocessing performed using *fMRIPprep* 1.2.3 (Esteban, Markiewicz, et al. (2018); Esteban, Blair, et al. (2018); RRID:SCR_016216), which is based on *Nipype* 1.1.6-dev (Gorgolewski et al. (2011); Gorgolewski et al. (2018); RRID:SCR_002502).

#### Anatomical data preprocessing

The T1-weighted (T1w) image was corrected for intensity non-uniformity (INU) using N4BiasFieldCorrection (Tustison et al. 2010, ANTs 2.2.0), and used as T1w-reference throughout the workflow. The T1w-reference was then skull-stripped using antsBrainExtraction.sh (ANTs 2.2.0), using OASIS as target template. Brain surfaces were reconstructed using recon-all (FreeSurfer 6.0.1, RRID:SCR_001847, Dale, Fischl, and Sereno 1999), and the brain mask estimated previously was refined with a custom variation of the method to reconcile ANTs-derived and FreeSurfer-derived segmentations of the cortical gray-matter of Mindboggle (RRID:SCR_002438, Klein et al. 2017). Spatial normalization to the ICBM 152 Nonlinear Asymmetrical template version 2009c (Fonov et al. 2009, RRID:SCR_008796) was performed through nonlinear registration with antsRegistration (ANTs 2.2.0, RRID:SCR_004757, Avants et al. 2008), using brain-extracted versions of both T1w volume and template. Brain tissue segmentation of cerebrospinal fluid (CSF), white-matter (WM) and gray-matter (GM) was performed on the brain-extracted T1w using fast (FSL 5.0.9, RRID:SCR_002823, Zhang, Brady, and Smith 2001).

#### Functional data preprocessing

For each of the 12 BOLD runs found per subject, the following preprocessing was performed. First, a reference volume and its skull-stripped version were generated using a custom methodology of *fMRIPrep*. A deformation field to correct for susceptibility distortions was estimated based on two echo-planar imaging (EPI) references with opposing phase-encoding directions, using 3dQwarp Cox and Hyde (1997) (AFNI 20160207). Based on the estimated susceptibility distortion, an unwarped BOLD reference was calculated for a more accurate co-registration with the anatomical reference. The BOLD reference was then co-registered to the T1w reference using bbregister (FreeSurfer) which implements boundary-based registration (Greve and Fischl 2009). Co-registration was configured with nine degrees of freedom to account for distortions remaining in the BOLD reference. Head-motion parameters with respect to the BOLD reference (transformation matrices, and six corresponding rotation and translation parameters) are estimated before any spatiotemporal filtering using mcflirt (FSL 5.0.9, Jenkinson et al. 2002). BOLD runs were slice-time corrected using 3dTshift from AFNI 20160207 (Cox and Hyde 1997, RRID:SCR_005927). The BOLD time-series (including slice-timing correction when applied) were resampled onto their original, native space by applying a single, composite transform to correct for head-motion and susceptibility distortions. These resampled BOLD time-series will be referred to as *preprocessed BOLD in original space*, or just *preprocessed BOLD*. The BOLD time-series were resampled to MNI152NLin2009cAsym standard space, generating a *preprocessed BOLD run in MNI152NLin2009cAsym space*. First, a reference volume and its skull-stripped version were generated using a custom methodology of *fMRIPrep*. Several confounding time-series were calculated based on the *preprocessed BOLD*: framewise displacement (FD), DVARS and three region-wise global signals. FD and DVARS are calculated for each functional run, both using their implementations in *Nipype* (following the definitions by Power et al. 2014). The three global signals are extracted within the CSF, the WM, and the whole-brain masks.

Additionally, a set of physiological regressors were extracted to allow for component-based noise correction (*CompCor*, Behzadi et al. 2007). Principal components are estimated after high-pass filtering the *preprocessed BOLD* time-series (using a discrete cosine filter with 128s cut-off) for the two *CompCor* variants: temporal (tCompCor) and anatomical (aCompCor). Six tCompCor components are then calculated from the top 5% variable voxels within a mask covering the subcortical regions. This subcortical mask is obtained by heavily eroding the brain mask, which ensures it does not include cortical GM regions. For aCompCor, six components are calculated within the intersection of the aforementioned mask and the union of CSF and WM masks calculated in T1w space, after their projection to the native space of each functional run (using the inverse BOLD-to-T1w transformation). The head-motion estimates calculated in the correction step were also placed within the corresponding confounds file. The BOLD time-series, were resampled to surfaces on the following spaces: *fsaverage6*. All resamplings can be performed with *a single interpolation step* by composing all the pertinent transformations (i.e. head-motion transform matrices, susceptibility distortion correction when available, and co-registrations to anatomical and template spaces). Gridded (volumetric) resamplings were performed using antsApplyTransforms (ANTs), configured with Lanczos interpolation to minimize the smoothing effects of other kernels (Lanczos 1964). Non-gridded (surface) resamplings were performed using mri_vol2surf (FreeSurfer).

After running fMRIprep (procedures described above), we additionally performed polynomial trend correction using AFNI’s 3dDeconvolve function and simultaneously regressed out six degrees of head motion (x, y, z, roll, pitch, yaw), first six aCompCor components, and associated cosine regressors. The first six volumes of each run were discarded to allow T1 equilibration.

Finally, we extracted ROIs in common space using the Destrieux (Destrieux et al., 2010) and Schaefer 400 parcellation 17 network atlases (Schaefer et al., 2018), resampled to 3.0 mm.

### Statistical analyses

We had two main approaches to investigate the impact of scramble level in various brain regions. First, to test the stimulus-driven reliability of neural responses in a given region (and thus, whether that region integrates context at the timescale corresponding to each scramble level), we used inter-subject correlation (ISC). This is similar to the logic of previous studies using scrambled narrative speech (Lerner et al, 2011). We then conducted repeated-measures ANOVAs to test interactions between feedback type (“AM” vs. “M” group), scramble level, and ROI. We also conducted bootstrapped permutation tests between individual scramble conditions. Second, to test the impact of scrambling on functional connectivity between regions, we computed intersubject functional correlation (ISFC), applying 2-way ANOVAs (between group and scramble level) for several ROI combinations, as well as bootstrapped permutation tests to contrast individual scramble conditions. Finally, we conducted several control analyses to test how reliably different brain regions represented musical content (“Inter-rep correlation” analyses below, by condition and by musical section), as well as controls to verify that observed effects of scrambling could not be explained by performers’ behavioral accuracy levels or lower-level cues in the stimuli (pitch height, rhythmic density). All analyses were computed using the averaged time series across all three (identical) runs (reps) of a given condition, with the last 10 TRs cropped, unless otherwise noted. We describe each analytical approach in further detail below.

#### Intersubject correlation (ISC)

To compute synchrony, we used ISC (Hasson et al., 2004; Lerner et al., 2011; Nastase et al., 2019), which isolates stimulus-driven activity that is reliable across participants while eliminating subject-specific intrinsic fluctuations and scanner noise. For each ROI, we computed Pearson correlations between each participant’s neural time series and the average time series across the remaining participants in the group (“AM”, “M”; Figure 2). We computed this ISC value for each participant in each of the scramble conditions, Fisher-z transformed these ISC values, and ran statistical tests on them.

First, we ran a three-way ANOVA including main effects of scramble condition, ROI, and group and their interaction in influencing ISC values. Next, in each ROI, we ran post-hoc contrasts comparing the Intact condition with the other three individual scramble conditions (all corrected using Bonferroni method, p < .005; see Table S1). Specifically, we ran bootstrapped permutation tests on ISC values by shuffling the labels across scrambling conditions and computing the mean difference (1000 iterations); then comparing the true mean difference to yield a 95% confidence interval.

#### Intersubject functional correlation (ISFC)

To measure how scrambling affected functional connectivity between regions, we computed intersubject functional correlation (ISFC; Simony et al., 2016). First, we tested the significance of ISFC across a set of core ROIs (Figs. 3, 4 & S3). Because of the presence of long-range temporal autocorrelation in the BOLD signal, we assessed the statistical likelihood of each observed ISFC value using a permutation (bootstrapping) procedure based on surrogate data generated using phase randomization (as in Simony et al., 2016). Phase-randomized surrogates have the same mean and autocorrelation as the original signal. We tested the null hypothesis that the BOLD signal (averaged across voxels within each ROI) in each individual was independent of the BOLD signal values in all other ROIs in any other individual at any point in time and corrected for multiple comparisons using the Benjamini-Hochberg procedure (q< .05). Next, we tested the systematic influence of scrambling on ISFC by conducting a series of two-way ANOVAs (group x scramble condition) for different groupings of ROIs. Finally, we conducted post-hoc bootstrapped permutation tests between individual scramble conditions (same procedure as for ISC; see above), with p < .007 after Bonferroni correction (see Table S4).

### Control analyses

#### Inter-rep correlation (IRC), by condition

To further confirm the stimulus-driven reliability of neural responses, we tested how well individual runs of a given scramble condition reflected musical content specific to that condition. Specifically, we computed Pearson correlations between repetitions (identical runs) of the same scramble condition, which we call “inter-rep correlation” (also known as “intrasubject correlation”; Hasson et al., 2009; Silbert et al., 2014), and compared them with correlations between different (randomly chosen) scrambled conditions (Figure S1). We conducted bootstrapped permutation tests between correlation values from real vs. shuffled (randomly chosen) conditions (p < .005 after Bonferroni correction; Table S2).

#### Inter-rep correlation (IRC), by musical section

In a second version of the inter-rep reliability test above, we computed IRC between repetitions of the Intact condition alone, comparing correlations between segments of music corresponding to either the same section of the piece or different (randomly chosen) sections of the piece (again using bootstrapped permutation tests; p < .005 after Bonferroni correction; Figure S2 and Table S3). Sections are depicted in color in Figure 1 (fourth column).

#### Behavioral accuracy

Performers’ behavioral accuracy was computed by comparing the MIDI output (recorded directly from the keyboard in GarageBand) to an ideal MIDI template (from the sheet music of each condition). For each beat, we computed the proportion of total correct MIDI values (including all pitch values that should have been released since the last beat as well as placed on the current beat) and averaged those across beats in each scramble condition for each participant (Figure S4).

#### Controlling for low-level musical cues

Finally, we tested the possibility that our reported effects of scramble condition might have been influenced by differences in lower-level musical cues across scramble conditions (specifically, changes in pitch height and rhythmic density that could have theoretically been relatively large after scramble boundaries). We measured a change in pitch height across a scrambled boundary as the difference between average pitch height (i.e., average MIDI value of all notes) in the beat following a scramble boundary vs. the beat preceding the boundary. We measured a change in rhythmic density as the difference between the rhythmic density (i.e., total number of note onsets) in the beat following a scramble boundary vs. the beat preceding the boundary. For each scramble condition (1B, 2B, 8B) we computed the changes in each of these two values across all scramble boundaries (Figure S5, orange boxes). Then, we extracted equivalently-sized segments of music in the Intact condition (i.e., one-bar, two-bar, and eight-bar segments of intact music; blue boxes) and conducted t-tests between the two types of segments.

## Acknowledgments

We thank Nicholas DePinto, Weiming Dai, Leigh Nystrom, and Mark Pinsk from the Scully Center at the Princeton Neuroscience Institute for their assistance in testing, troubleshooting, and setting up the scanner environment to be compatible with our MR-safe keyboard. We thank Jacky Lu for manufacturing the keyboard. We thank Madeline Kushan for assistance with stimulus development, recruitment, and piloting. We thank Samuel Nastase for leading a community for building and sharing tools to enhance fMRI preprocessing at Princeton. We also thank Uri Hasson and other members of the Hasson Lab at Princeton and SoNIC Lab at the University of Rochester for guidance and feedback. This work was supported by a GRAMMY MuseumⓇ Research Grant awarded to E.A.P., C.R.I., and Uri Hasson, a C.V. Starr Fellowship awarded to E.A.P., a F99 NS118740 awarded to J.W., and R01 MH112357 awarded to Uri Hasson.

## Author contributions

E.A.P., J.W., and C.R.I. jointly conceptualized the study and designed the methodology.

E.A.P. led the data collection, as well as the data curation, analysis, and visualization.

C.R.I. and J.W. contributed substantially to data curation, analysis, and visualization. R. C-C. contributed to data curation and formal analysis. E.A.P. supervised the project and wrote the original draft of the manuscript, and J.W., C.R.I., and R. C-C. contributed to manuscript review and editing. J.W. and C.R.I. contributed equally.

