## Supplementary Information for "Hierarchical neural integration of musical structure during expert performance"

| **"AM"** | A1 | STG | V1 | PMd | PMv | Motor Cortex | preSMA | SMA | Cerebellum | DMN | Hippocampus |
|---|---|---|---|---|---|---|---|---|---|---|---|
| I vs. 1B | .17 | .049 | <.001 | <.001 | <.001 | <.001 | .18 | <.001 | .18 | .31 | <.001 |
| I vs. 2B | <.001 | .05 | <.001 | <.001 | <.001 | <.001 | <.001 | <.001 | <.001 | <.001 | <.001 |
| I vs. 8B | .29 | .28 | .05 | .41 | .19 | .12 | .51 | .32 | <.001 | <.001 | .11 |

| **"M"** | A1 | STG | V1 | PMd | PMv | Motor Cortex | preSMA | SMA | Cerebellum | DMN | Hippocampus |
|---|---|---|---|---|---|---|---|---|---|---|---|
| I vs. 1B | .63 | .70 | .19 | .26 | .09 | .20 | .39 | .34 | .09 | .90 | .93 |
| I vs. 2B | .54 | .36 | <.001 | .41 | .15 | .44 | .59 | .39 | <.001 | .58 | <.001 |
| I vs. 8B | .96 | .99 | .07 | .92 | .95 | .78 | .80 | .84 | .02 | .92 | .41 |

**Table S1**. P-values from permutation tests contrasting intersubject correlation (ISC) between scramble conditions (Intact vs. 1B, Intact vs. 2B, Intact vs. 8B) in "AM" (top) and "M" (bottom). $p < .005$ (.05/11) after Bonferroni correction.

Representations of fine-grained musical content during live performance

Because our stimuli contained detailed musical content (e.g., sections containing distinct themes in the Intact piece; segments of tonally intact musical structure that varied in size across scramble conditions), we exploited this rich structure to verify the reliability of music-related patterns in the neural data. Specifically, we tested how well different brain regions represented this content. Here, we took advantage of the fact that each subject played three repetitions of each scramble condition (see Methods for details). First, we tested whether we could classify the scramble condition that the subjects were playing by correlating repetitions of the same condition (which we call "inter-rep correlation", or IRC) and comparing these values to correlations between repetitions of different (randomly chosen) conditions, using bootstrapped permutation tests. In the vast majority of ROIs, in both the "AM" and "M" group, repetitions of the same scramble condition (e.g., "2B" vs. "2B"; green boxes in Figure S1) were more strongly correlated than repetitions of different scramble conditions ("2B" vs. "8B"; gray boxes). See Table S2 for full set of results (bootstrapped permutation tests; $p < .005$ after Bonferroni correction).

Second, we tested whether we could classify which *section* of the Intact Tchaikovsky piece subjects were performing at a given time (see Figure 1 for a schematic of the musical structure of the Intact condition; colors indicate sections, which featured distinct but cohesive melodic themes, each in a different key). Here, we computed inter-rep correlation (IRC) between repetitions of the Intact condition, comparing correlations between segments of music corresponding to either the same section of the piece or different (randomly chosen) sections of the piece, again using bootstrapped permutation tests. Again, in almost all ROIs, in both the "AM" and "M" group, repetitions of the same musical section (green boxes in Figure S2) were more strongly correlated than repetitions of different sections (gray boxes). See Table S3 for full set of results (bootstrapped permutation tests; $p < .005$ after Bonferroni correction).

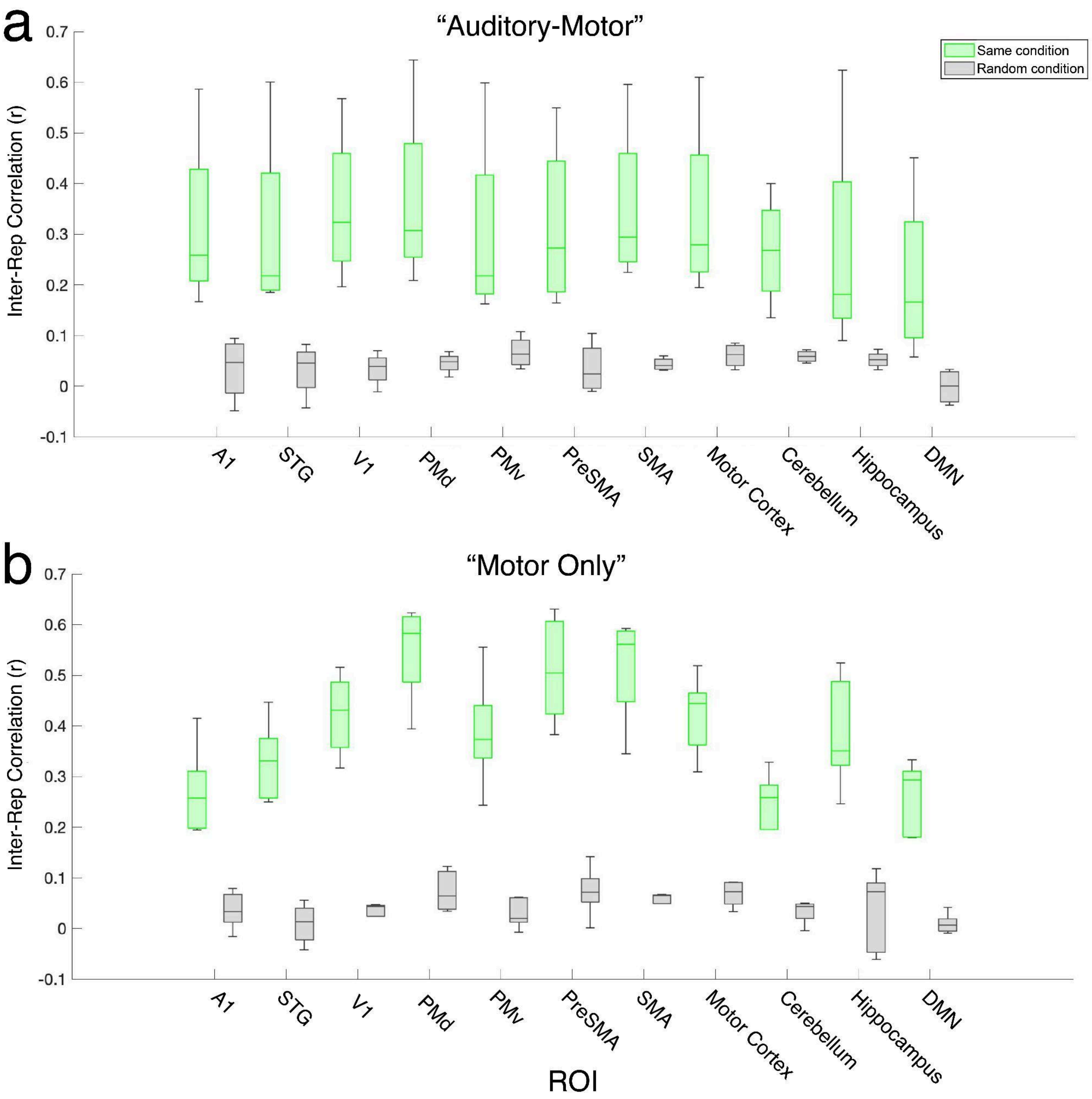


**Figure S1**. Within-subject reliability of responses (measured as inter-repetition correlation, or IRC), between repetitions of the same scramble condition (green) vs. randomly-chosen conditions (gray) in the (a) "AM" group and (b) "M" group. Boxcharts depict medians, with edges = 25th/75th percentile.

| | A1 | STG | V1 | PMd | PMv | PreSMA | SMA | Motor Cortex | Cerebellum | Hippocampus | DMN |
|---|---|---|---|---|---|---|---|---|---|---|---|
| "AM" | <.001 | <.001 | <.001 | <.001 | <.001 | <.001 | <.001 | <.001 | <.001 | <.001 | <.001 |
| "M" | <.001 | <.001 | <.001 | <.001 | <.001 | <.001 | <.001 | <.001 | <.001 | <.001 | <.001 |

**Table S2**. P-values from permutation tests contrasting inter-rep correlation (IRC) between repetitions of the same scramble condition (e.g., 2B vs. 2B) vs. different scramble conditions (e.g., 2B vs. 8B). p < .005 (.05/11) after Bonferroni correction.

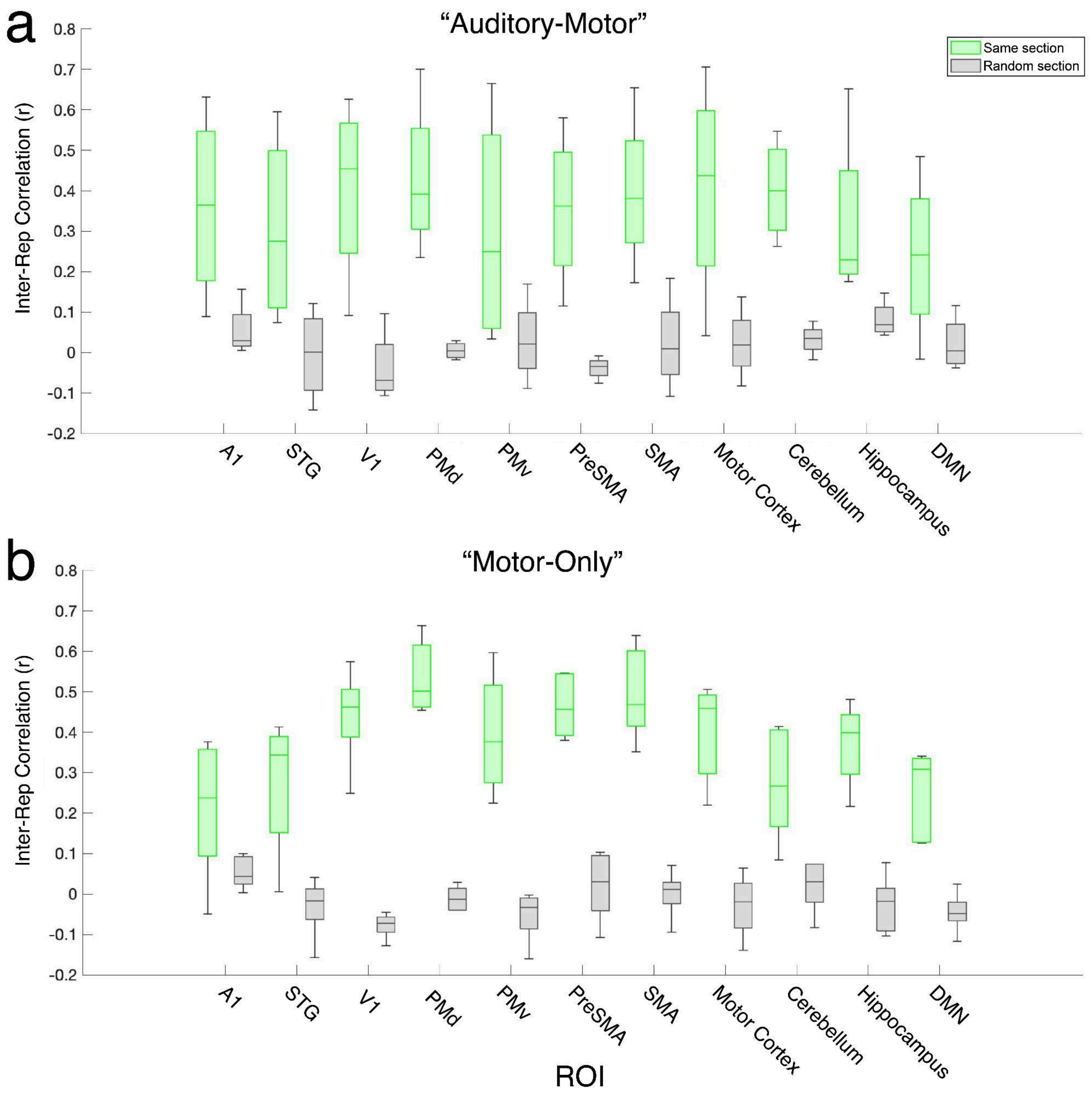


**Figure S2**. Within-subject response reliability (inter-repetition correlation, or IRC) between repetitions of the same section of music (green) vs. randomly-chosen sections (gray) in the (a) "AM" group and (b) "M" group. Boxcharts depict medians, with edges = 25th/75th percentile.

| | A1 | STG | V1 | PMd | PMv | PreSMA | SMA | Motor Cortex | Cerebellum | Hippocampus | DMN |
|---|---|---|---|---|---|---|---|---|---|---|---|
| "AM" | .06 | <.001 | <.001 | <.001 | .12 | <.001 | <.001 | .07 | <.001 | <.001 | .05 |
| "M" | .07 | .03 | <.001 | <.001 | <.001 | <.001 | <.001 | <.001 | <.001 | <.001 | <.001 |

**Table S3**. P-values from permutation tests contrasting inter-rep correlation (IRC) between the same section (theme) of music vs. different sections. $p < .005$ (.05/11) after Bonferroni correction.

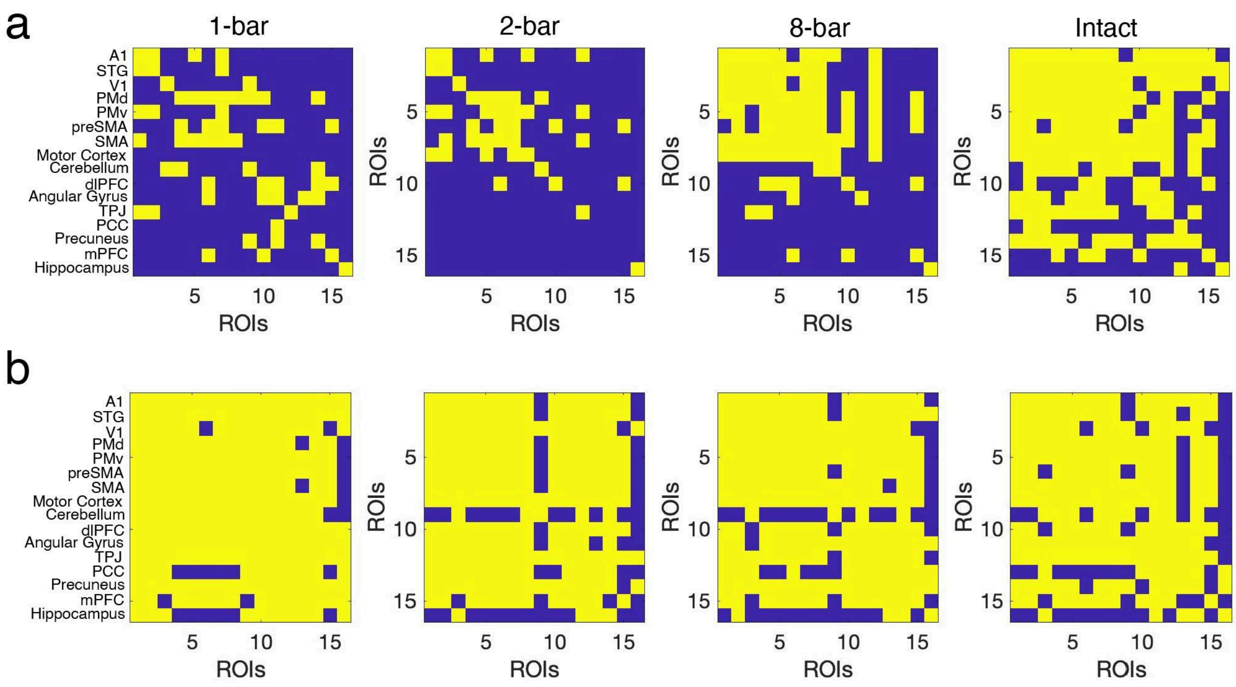


**Figure S3**. Significance values (yellow) for intersubject functional correlation (ISFC) analysis in (a) "AM" subjects and (b) "M" subjects (raw ISFC values depicted in Fig. 3). Significance was determined using a permutation procedure based on phase-randomized surrogate data (as in Simony et al., 2016) and corrected for multiple comparisons with the Benjamini-Hochberg false-discovery-rate procedure (q<.05).

| **"AM"** | A1-to-V1 | Auditory-motor | Auditory-DMN | Within-Motor | Motor-DMN | Cerebellum-to-DMN | V1-to-DMN |
|---|---|---|---|---|---|---|---|
| I vs. 1B | <.001 | .06 | .05 | <.001 | <.001 | <.001 | <.001 |
| I vs. 2B | .06 | .06 | <.001 | <.001 | <.001 | <.001 | <.001 |
| I vs. 8B | .17 | .26 | .05 | .25 | <.001 | <.001 | <.001 |

| **"M"** | A1-to-V1 | Auditory/ motor | Auditory-DMN | Within-Motor | Motor-DMN | Cerebellum-to-DMN | V1-to-DMN |
|---|---|---|---|---|---|---|---|
| I vs. 1B | .40 | .06 | .72 | .29 | .58 | .73 | .71 |
| I vs. 2B | .68 | .43 | .62 | .37 | .85 | .06 | .85 |
| I vs. 8B | .25 | .98 | .95 | .95 | .94 | .17 | .13 |

**Table S4**. P-values from permutation tests contrasting intersubject functional correlation (ISFC) between scramble conditions (Intact vs. 1B, Intact vs. 2B, Intact vs. 8B) in "AM" (top) and "M" (bottom) groups. p < .007 (.05/7) after Bonferroni correction.

Effects of scrambling cannot be explained by differences in behavioral accuracy/performance

Mean performance accuracy does not differ significantly across conditions (Kruskal-Wallis tests, “AM” group: $\chi^2(3) = 1.39$, $p = .71$; “M” group: $\chi^2(3) = 2.41$, $p = .49$), so this cannot explain large scrambling effects (e.g, ISC in motor cortex). In addition, different specific patterns of ISC and ISFC between the “AM” and “M” groups cannot be explained by differences in performance accuracy across the groups, since they did not perform significantly differently in any scramble condition (1B: $t(7) = -.45$, $p = .66$; 2B: $t(7) = -.61$, $p = .56$; 8B: $t(7) = .02$, $p = .99$; I: $t(7) = -.29$, $p = .78$).

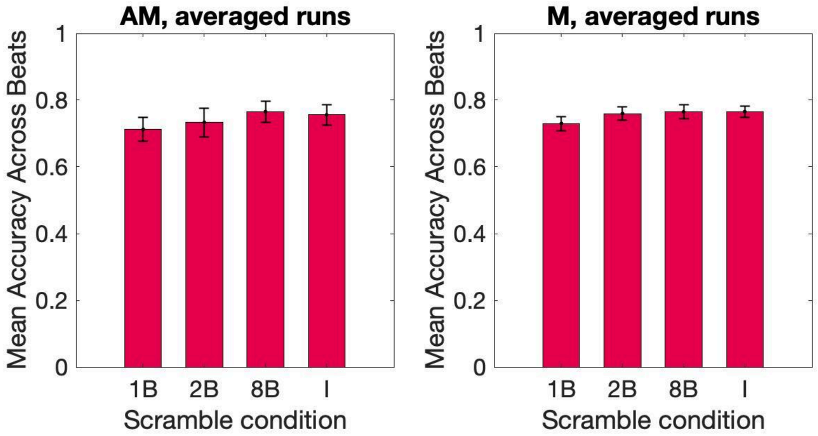


**Figure S4**. Mean performance (piano-playing) accuracy, averaged across all beats in a run of music, for each scramble condition in the “AM” (left) and “M” (right) groups. Error bars are s.e.m.

Impact of scrambling cannot be explained by relatively low-level musical cues

Changes in pitch height (Figure S5a) and rhythmic density (Figure S5b) across scramble boundaries (in the 1B, 2B, and 8B conditions; orange boxes) are not significantly greater than changes in these features across boundaries separating equivalently-sized segments of music in the Intact condition (i.e., one-bar, two-bar, and eight-bar chunks of intact music; blue boxes). T-tests for pitch height, scrambled condition vs. equivalent chunk of Intact music: one-bar ($p = .83$), two-bar ($p = .96$), eight-bar ($p = .72$). T-tests for rhythmic density: one-bar ($p = .76$), two-bar ($p = .89$), eight-bar ($p = .83$).

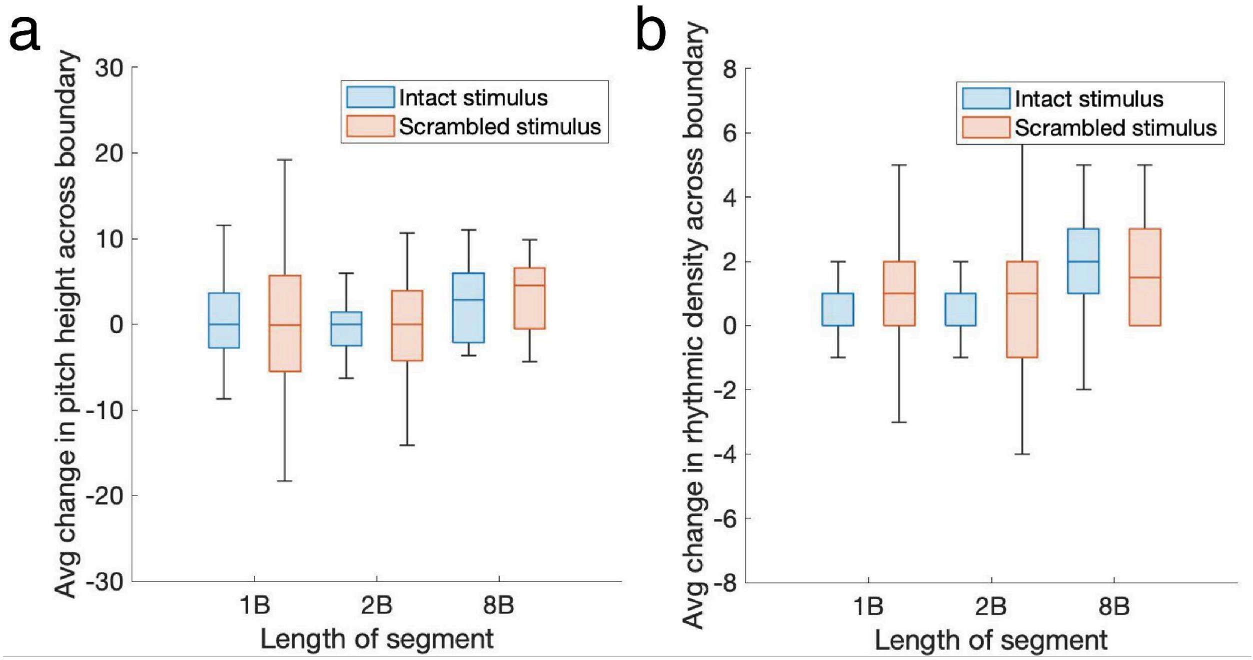


**Figure S5**. Analysis of low-level musical cues in our stimulus set. Average change in pitch height (left) and rhythmic density (right) across scramble boundaries for each scramble condition (orange boxes) and across boundaries separating equivalently-sized segments of music in the Intact condition (blue boxes). Boxcharts depict medians, with edges = 25th/75th percentile.

Head motion is minimal, despite the unique demands of the task

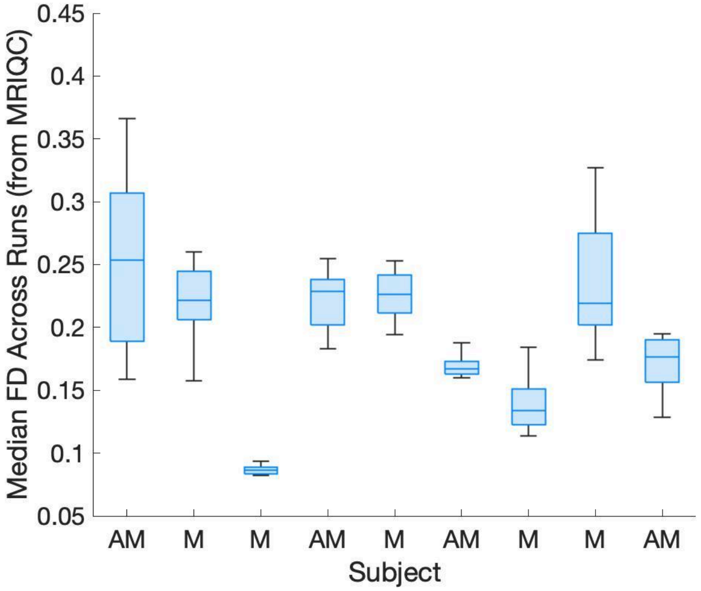


**Figure S6**. Median framewise displacement (FD), in mm, across all runs in each subject (MRIQC output from fMRIprep). Box edges = 25th/75th percentile.
